# c-MYC is an aggregation-prone, amyloidogenic protein

**DOI:** 10.64898/2026.03.12.711438

**Authors:** Ling Lin, Kun-Han Chuang, Chengkai Dai

**Affiliations:** Mouse Cancer Genetics Program, Center for Cancer Research, National Cancer Institute, Frederick, MD 21702, USA; School of Basic Medical Sciences, Fujian Medical University, Fuzhou, Fujian 350108, China

## Abstract

The transcription factor c-MYC is a central regulator of diverse biological processes and an oncogenic driver. Paradoxically, c-MYC also exhibits intrinsic tumor-suppressive activity through apoptosis, a phenomenon known as the “c-MYC paradox”. Here, we identify c-MYC as an intrinsically amyloidogenic protein. Under proteotoxic stress, c-MYC becomes detergent-insoluble aggregates. Notably, even in the absence of stress, c-MYC assembles into soluble amyloid-like oligomers in human cancer tissues and Alzheimer’s disease brains. *In vitro*, recombinant c-MYC proteins spontaneously form amyloid oligomers and protofibrils. In contrast, its obligate dimerization partner MAX is non-amyloidogenic and suppresses c-MYC amyloidogenesis. Mapping studies identify two short linear sequences within intrinsically disordered regions that confer amyloidogenicity. Importantly, the amyloid state of c-MYC induces apoptosis largely independently of transcription. Thus, these findings reveal a previously unrecognized amyloidogenic property of c-MYC, which contributes to its tumor-suppressing activity. Conceptually, c-MYC amyloidogenesis may serve as a built-in failsafe mechanism to restrain its oncogenic potential.

## INTRODUCTION

The bHLH/ZIP transcription factor c-MYC plays vital roles in human biology, functioning as both a potent oncogenic driver and a pluripotency-reprogramming factor.^1,2^ By heterodimerizing with MYC-associated factor X (MAX), MYC/MAX complexes bind the canonical E-box element (5’-CACGTG-3’) and its variants to regulate approximately 10-15% of the human genome.^1,3^ Dimerization with MAX is widely regarded as a prerequisite for c-MYC transcriptional activity.^4^ In normal, non-transformed cells, c-MYC undergoes rapid turnover and is maintained at low levels; in cancer, however, its expression is frequently elevated through gene amplification, chromosomal translocation, or increased protein stability.^5^ Despite its prominent oncogenic role, c-MYC paradoxically induces apoptosis—an intrinsic tumor-suppressive mechanism—particularly under conditions of cellular stress or elevated expression, an activity largely attributed to its transcriptional function.^6–9^

Proteins adopt functional conformations in their soluble, properly folded states; however, proteotoxic stress and genetic mutations can drive protein misfolding and aggregation, causing both loss of function and cytotoxicity. Amyloids are a distinct class of protein aggregates characterized by cross-β-sheet structures and are strongly implicated in human diseases, especially neurodegenerative disorders.^10^ Two fluorescent dyes—Congo red (CR) and thioflavin T (ThT)—are widely used to detect amyloids through direct binding.^11^ In addition, the conformation-specific antibody A11 serves as a valuable tool for recognizing soluble prefibrillar amyloid oligomers (AOs), a highly neurotoxic conformer whose recognition is independent of primary amino acid sequences.^12^ Beyond its diagnostic utility, CR also inhibits amyloid formation.^13,14^

Here, we report that c-MYC, but not its obligate dimerization partner MAX, forms detergent-insoluble aggregates upon heat shock (HS). Furthermore, in human cancer tissues and, unexpectedly, in Alzheimer’s disease (AD) brains, c-MYC assembles into detergent-soluble prefibrillar oligomers detectable by the A11 antibody even in the absence of HS. Notably, both aggregation and oligomerization are suppressed by CR. *In vitro*, recombinant c-MYC proteins spontaneously form AOs and protofibrils, demonstrating its intrinsic amyloidogenic capacity, whereas MAX both fails to form amyloids and actively suppresses c-MYC amyloidogenesis. Systematic screening of a synthetic peptide library derived from human c-MYC identified two intrinsically disordered short linear sequences (aa20-39 and aa210-228) as amyloidogenic determinants. Alanine-scanning mutagenesis further revealed that a large proportion of residues within both fragments contribute to amyloid formation. Strikingly, a transcription-deficient c-MYC mutant lacking the C-terminal MAX-dimerization and DNA-binding domain retains the ability to induce apoptosis, and deletion of the two amyloidogenic regions markedly attenuates this activity. Together, these findings establish c-MYC as an aggregation-prone, intrinsically amyloidogenic protein whose amyloid state promotes apoptosis independently of transcription, revealing a previously unrecognized mechanism underlying its tumor-suppressive function.

These results provide new insights into the longstanding c-MYC paradox and reveal its complex implications in human pathologies, including cancer and neurodegeneration.

## RESULTS

### c-MYC acquires amyloid-like properties under stress and pathological conditions

We recently showed that heat shock factor 1 (HSF1) potentiates c-MYC-mediated transcription through physical interactions under non-stressed conditions, representing a non-canonical transcriptional action of HSF1.^15^ We therefore examined how this c-MYC−HSF1 interaction is affected by proteotoxic stress—specifically HS—during which HSF1 is recruited to orchestrate the canonical heat-shock, or proteotoxic stress, response (HSR/PSR).^16,17^

In HeLa cells, transient HS caused a marked depletion of detergent-soluble c-MYC, accompanied by a corresponding accumulation of detergent-insoluble c-MYC (Figure 1A), indicative of stress-induced protein aggregation. In contrast, its obligate dimerization partner MAX and the major cellular chaperones HSP90 and HSC/HSP70 remained in the soluble fraction. Importantly, CR partially blocked c-MYC aggregation (Figure 1A), consistent with its known capacity to inhibit amyloidogenesis, though broader effects on protein aggregation cannot be excluded.

**Figure 1:**
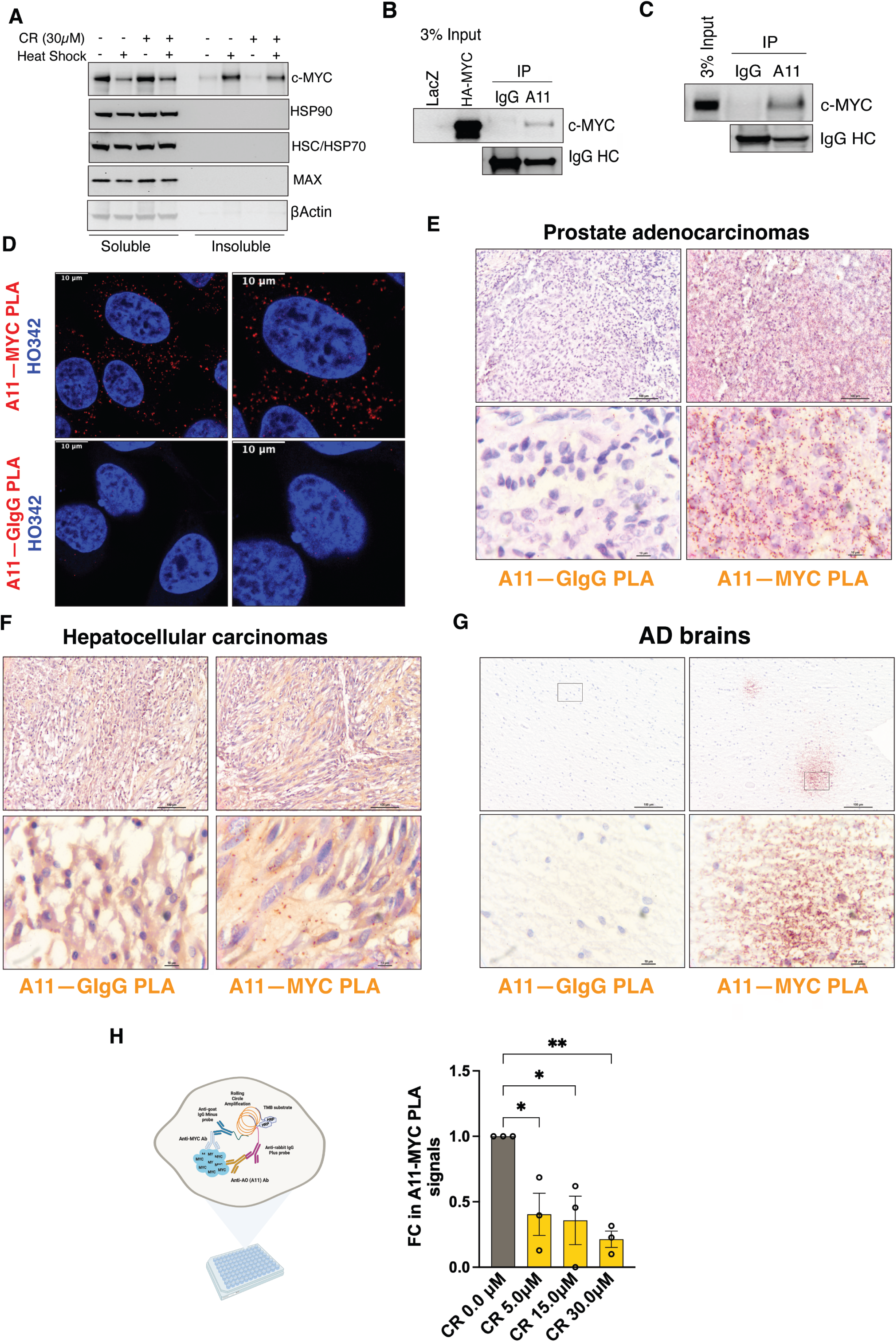
c-MYC becomes detergent-insoluble and displays amyloid-like properties under proteotoxic stress and pathological conditions. (A) Detection of detergent-soluble and - insoluble c-MYC proteins by immunoblotting in HeLa cells pre-treated with 30 μM CR, followed by HS at 43°C for 30 min (representative images of three independent experiments). (B) and (C) Immunoprecipitation of either exogenously expressed HA-c-MYC or endogenous c-MYC proteins in HeLa cells by A11 antibodies (representative images of three independent experiments). (D) Detection of endogenous c-MYC proteins in HeLa cells by PLA using both the goat anti-c-MYC Ab and the rabbit anti-AO (A11) Ab. Nuclei are counterstained with Hoechst 33342. Scale bar: 10 μm. (E)-(G) Detection of endogenous c-MYC proteins in human prostate adenocarcinoma, hepatocellular carcinoma, and Alzheimer’s disease brain tissues by brightfield PLA using both the goat anti-c-MYC Ab and the rabbit anti-AO (A11) Ab. Scale bar: 100 μm for low magnification and 10 μm for high magnification. (H): Quantitation of endogenous c-MYC recognized by both the goat anti-c-MYC Ab and the rabbit anti-AO (A11) Ab through In-Cell PLA ELISA in HeLa cells treated with CR (mean ± SD, n=3 independent experiments, One-way ANOVA).

To determine whether the aggregates are amyloid-like, we employed the conformation-specific A11 antibody, which has been widely applied to recognize soluble prefibrillar AOs in a sequence-independent manner.^12^ A11 immunoprecipitation captured both exogenous HA-c-MYC and endogenous c-MYC proteins in HeLa cells even without HS (Figures 1B and 1C). These findings were further corroborated by *in situ* Proximity Ligation Assay (PLA), ^18^ which detects two distinct epitopes within the same protein or protein-protein interactions (Figure 1D). The PLA signals were quantified by flow cytometry (Figure S1A). Notably, these PLA signals were predominantly cytoplasmic (Figure 1D), suggesting that these c-MYC conformers are transcriptionally inactive. These observations were not an artifact of cell culture, as A11^+^-c-MYC oligomers were readily detected in both primary and xenografted human cancer tissues (Figures 1E, 1F, and S1B). Strikingly, analogous oligomers were identified in human AD brain tissues (Figure 1G), where they resembled diffuse Aβ plaques. CR treatment suppressed A11^+^-c-MYC oligomer formation (Figures 1H and S1C), reinforcing their amyloidogenic nature. Collectively, these results demonstrate that c-MYC acquires amyloid-like properties both under proteotoxic stress and in pathological settings.

### c-MYC is intrinsically amyloidogenic

Because immunoprecipitation and PLA cannot discriminate between c-MYC interacting with pre-existing AOs and c-MYC itself adopting an amyloid conformation, we turned to *in vitro* assays using purified recombinant proteins.

In classical fibrillation assays, recombinant c-MYC proteins from two independent sources spontaneously assembled into amyloid fibrils, as measured by ThT fluorescence (Figure 2A). This contrasted sharply with GST and a panel of transcription factors, including HSF1, USF1, OCT4, and MAX, none of which showed fibrillation. Notably, neither MAX, the obligate dimerization partner of c-MYC, nor USF1, another member of the c-MYC family, formed amyloid fibrils, underscoring the specificity of c-MYC amyloidogenicity. Beyond fibrils, recombinant c-MYC also formed soluble A11^+^–Aos (Figure 2B). GST-c-MYC proteins produced substantially higher AO levels, likely reflecting the solubilizing effect of the GST tag. Transmission electron microscopy (TEM) directly confirmed the assembly of c-MYC protofibrils (Figure 2C). Collectively, these *in vitro* data establish that c-MYC is intrinsically amyloidogenic.

**Figure 2:**
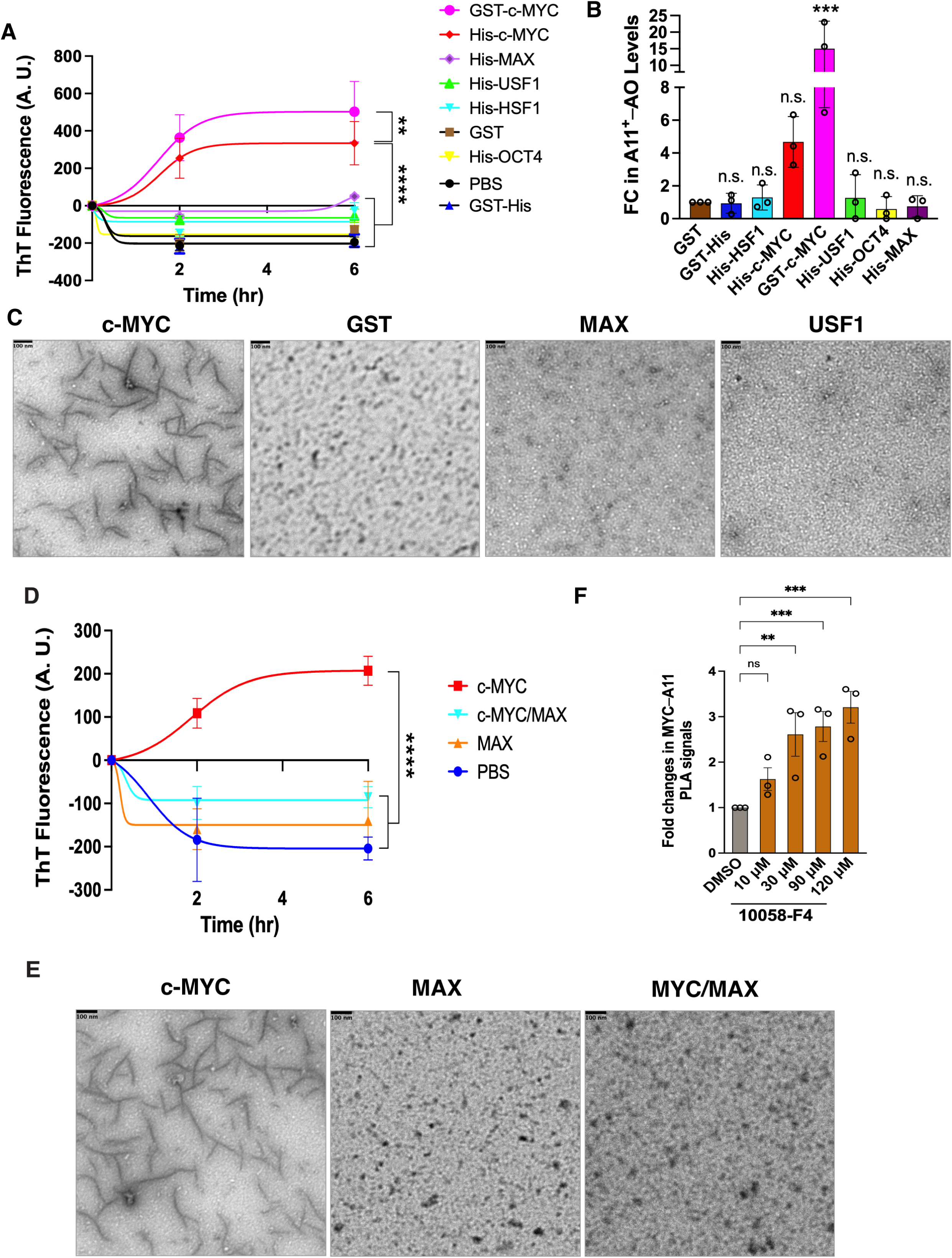
c-MYC is intrinsically amyloidogenic. (A) *In vitro* fibrillation assays using recombinant c-MYC proteins (mean ± SEM, n=3 independent experiments, Two-way ANOVA). (B) ELISA quantitation of A11^+^−AOs generated by recombinant c-MYC proteins *in vitro* (mean ± SD, n=3 independent experiments, One-way ANOVA). (C) TEM visualization of protofibrils formed by recombinant c-MYC proteins *in vitro* (representative images of three independent experiments). Scale bars: 100 nm. (D) *In vitro* fibrillation assays using recombinant c-MYC /MAX dimers (mean ± SEM, n=3 independent experiments, Two-way ANOVA). (E) TEM visualization of recombinant c-MYC/MAX dimers *in vitro* (representative images of three independent experiments). Scale bars: 100 nm. (F) Quantitation of endogenous c-MYC AOs in HeLa cells treated with 10058-F4 by In-Cell PLA ELISA (mean ± SD, n=3 independent experiments, One-way ANOVA).

Given that c-MYC physiologically dimerizes with non-amyloidogenic protein MAX,^4^ we next asked whether this interaction modulates c-MYC amyloidogenesis. Recombinant c-MYC/MAX heterodimers failed to form amyloid fibrils, in stark contrast to c-MYC alone (Figures 2D and 2E). Consistently, disrupting MYC/MAX dimerization in HeLa cells with the small molecule 10058-F4 increased A11^+^–c-MYC oligomers, despite reducing detergent-soluble c-MYC (Figures 2F, S2A, and S2B).^19^ These results indicate that dimerization with MAX actively suppresses c-MYC amyloid formation.

### Mapping the intrinsic amyloidogenic regions of c-MYC

To systematically identify amyloidogenic regions, we designed a library of 24 synthetic peptides spanning the human c-MYC protein, each comprising 19 non-overlapping amino acids— an approach previously validated for mapping protein-protein interaction interfaces.^15^ In vitro fibrillation assays identified two peptides—P2 (aa20-38) and P12 (aa210-228)—as amyloidogenic, mirroring the behavior of full-length c-MYC (Figure 3A). Of interest, only P12 consistently generated A11^+^–AOs (Figure 3B). Both peptides reside within intrinsically disordered regions (IDRs) of c-MYC (Figure S3A).

**Figure 3:**
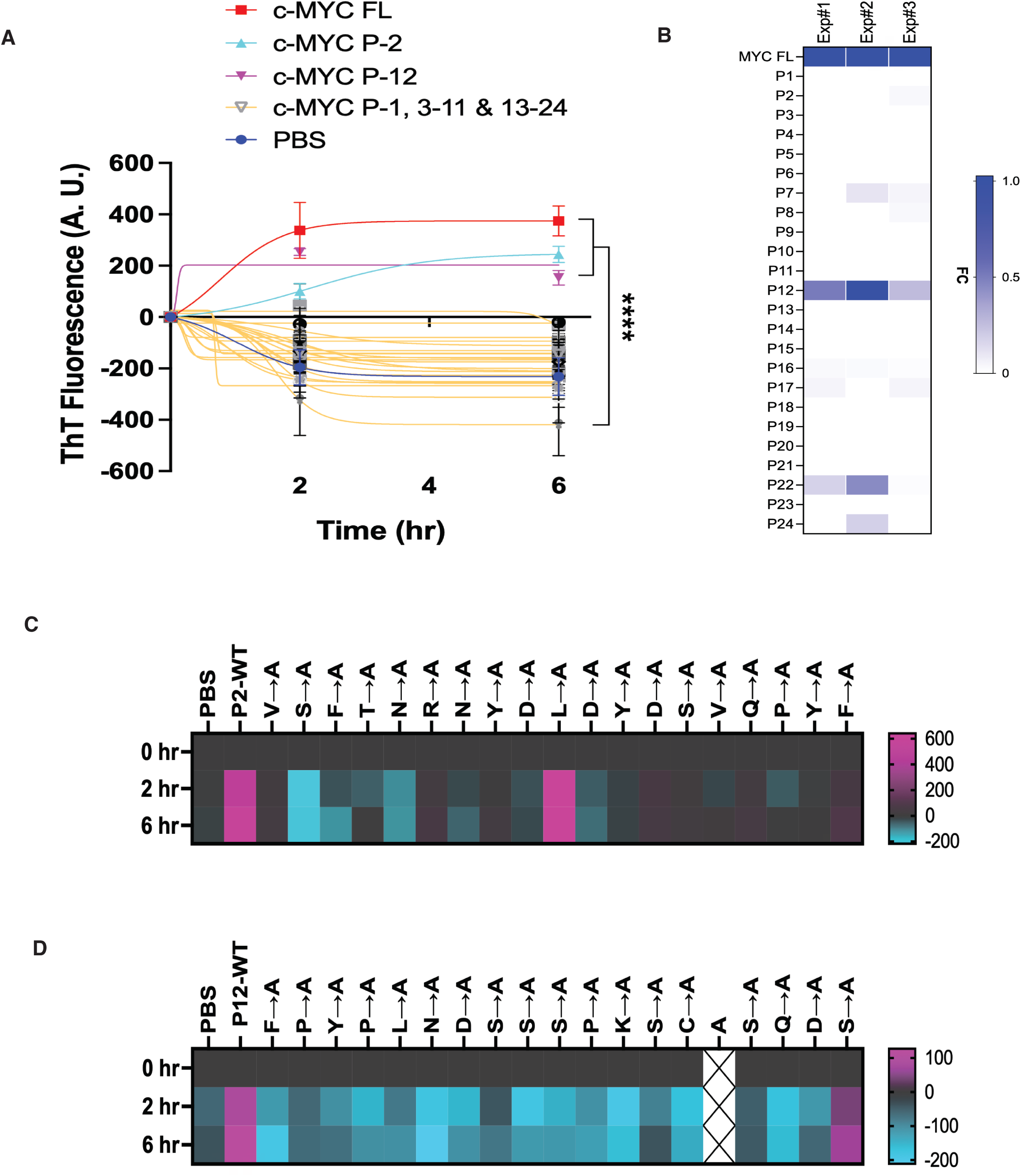
Decoding the amyloidogenicity of c-MYC *in vitro*. (A) *In vitro* fibrillation assays using the human c-MYC peptide library (mean ± SEM, n=3 independent experiments, Two-way ANOVA). (B) ELISA quantitation of A11^+^−AOs generated by synthetic c-MYC peptides. Results of three independent experiments are presented as a heatmap. FC: fold change. (C) and (D) *In vitro* fibrillation assays using the P2 and P12 peptides that are subjected to alanine scan. Results are presented as heatmaps using the averages of three independent experiments.

To resolve the individual residues responsible for amyloid formation, we subjected P2 and P12 to alanine-scanning mutagenesis. Single-residue alanine substitution generally caused marked reductions in amyloid fibril formation for both peptides (Figures 3C and 3D), demonstrating that multiple residues within each intrinsically disordered sequence collectively govern c-MYC amyloidogenicity.

### Amyloidogenesis drives transcription-independent apoptosis

Amyloid conformers are frequently cytotoxic, leading us to hypothesize that c-MYC amyloidogenesis contributes to apoptosis and thereby underlies its tumor-suppressive activity. It is well established that c-MYC induces apoptosis under stress conditions, including serum starvation and DNA damage, or upon elevated expression. ^6–9^ To isolate amyloid-dependent apoptosis from transcription-dependent apoptosis, we generate a transcription-deficient mutant, HA-c-MYC^ΔC^, lacking the C-terminal DNA-binding and MAX-dimerization domains while retaining the amyloidogenic regions (Figure 4A). In serum-starved NIH-3T3 cells, overexpression of HA-c-MYC^WT^ induced apoptosis, as measured by caspase 3 cleavage (Figure 4B). Strikingly, so did HA-c-MYC^ΔC^ mutants (Figures 4B), suggesting the existence of a transcription-independent mechanism.

**Figure 4:**
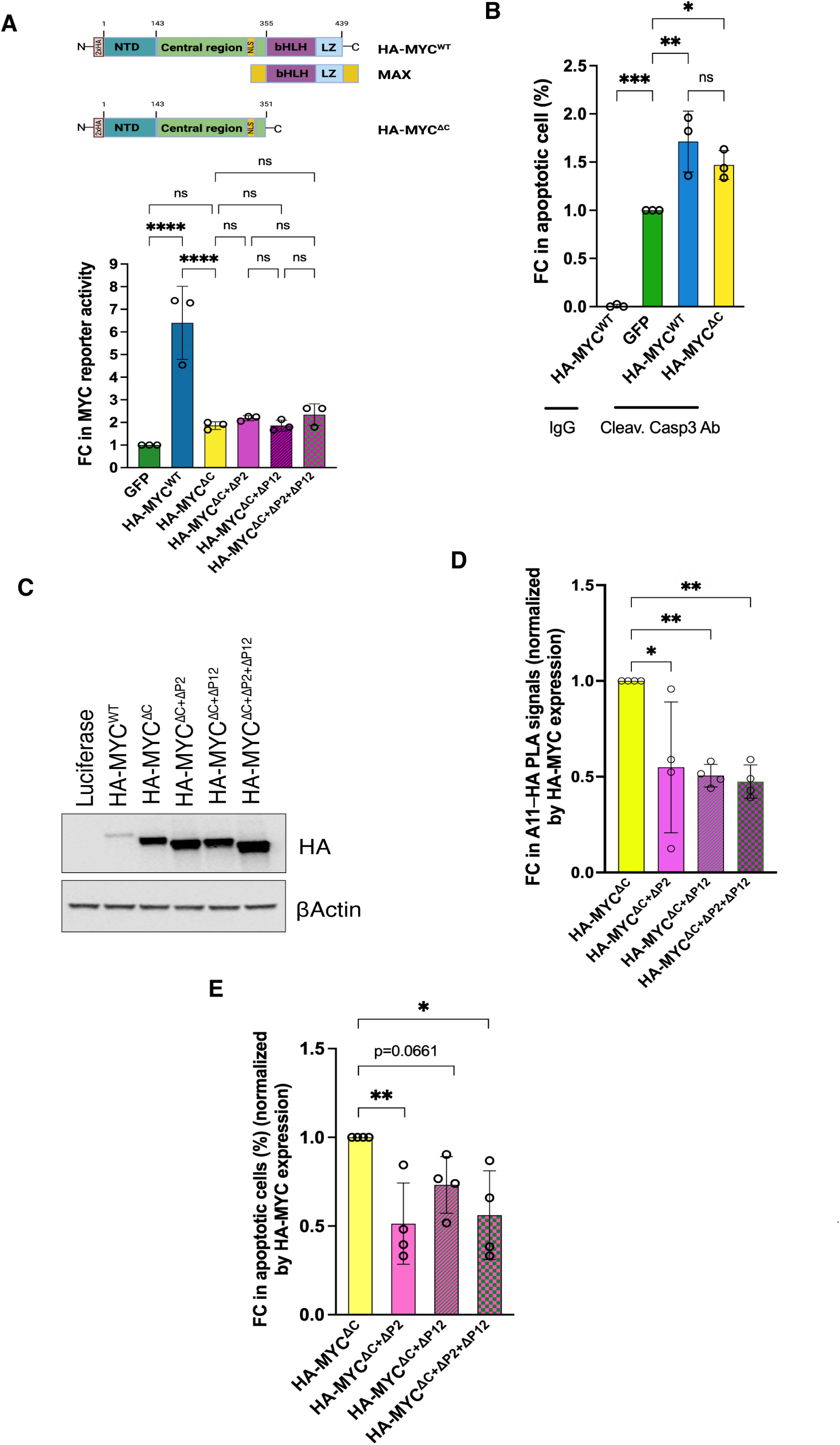
The amyloidogenesis of c-MYC contributes to its intrinsic tumor suppressor activity. (A) Measurements of the transcriptional activities of HA-MYC^WT^ and HA-MYC^ΔC^ in HeLa cells using the dual MYC reporter system (mean ± SD, n=3 independent experiments, One-way ANOVA). (B) Quantitation of apoptosis induced by transient expression of HA-MYC^WT^ or HA-MYC^ΔC^ in serum-starved NIH3T3 cells by flow cytometry using anti-cleaved caspase 3 Abs (mean ± SD, n=3 independent experiments, One-way ANOVA). (C) Detection of HA-MYC expression in serum-starved NIH3T3 cells by immunoblotting (representative images of four independent experiments). (D) Quantitation of A11^+^–HA-MYC^ΔC^ and its deletion mutants in NIH3T3 cells by In-Cell PLA ELISA using rabbit A11 Abs and mouse anti-HA Abs. The results are normalized by HA-MYC expression levels (mean ± SD, n=4 independent experiments, One-way ANOVA). (E) Quantitation of apoptosis induced by transient expression of HA-MYC^ΔC^ and its deletion mutants in serum-starved NIH3T3 cells by flow cytometry using anti-cleaved caspase 3 Abs. The results are normalized by HA-MYC expression levels (mean ± SD, n=4 independent experiments, One-way ANOVA).

To determine the contribution of amyloidogenesis, we deleted the P2 and P12 regions from HA-c-MYC^ΔC^. As deletions affected protein expression levels (Figure 4C), apoptotic readouts were normalized accordingly. Deletion of the P2 region substantially reduced apoptosis, whereas deletion of the P12 region had a more modest effect (Figure 4D). Critically, removal of these amyloidogenic regions did not further impair the residual transcriptional activity of HA-c-MYC^ΔC^ mutants (Figure 4A), yet significantly diminished its amyloidogenic and pro-apoptotic capacities (Figures 4D and 4E), thereby decoupling apoptosis induction from transcription. In aggregate, these results establish that c-MYC amyloidogenesis is a driver of cytotoxicity and contributes to its intrinsic tumor suppressor function.

## DISCUSSION

### **c-** MYC is an aggregation-prone and amyloidogenic protein

Our combined *ex vivo* and *in vitro* analyses provide convergent evidence that c-MYC is an aggregation-prone, intrinsically amyloidogenic protein. This conclusion is supported by multiple independent lines of evidence: CR-mediated inhibition of c-MYC aggregation and oligomerization; ThT binding indicative of c-MYC amyloid fibril formation; recognition of c-MYC AOs by the conformation-specific A11 antibody; and direct visualization of c-MYC protofibrils by TEM. Importantly, we identified two intrinsically disordered short linear sequences as the primary determinants of c-MYC amyloidogenicity. Although both regions assembled into amyloid fibrils *in vitro*, as evidenced by ThT binding, only P12 consistently produced A11^+^–AOs, raising the possibility that P2 generates a conformationally distinct class of oligomers not recognized by A11.

### MAX suppresses c-MYC amyloidogenesis

A central finding of this study is that MAX—itself non-amyloidogenic—suppresses c-MYC amyloid formation. This observation has important physiological implications: in primary non-transformed cells, where c-MYC expression is low and MAX is sufficiently available for dimerization, c-MYC amyloidogenesis is effectively restrained.

In cancer, this balance is disrupted. *c-MYC* is frequently overexpressed, yet its expression levels are poorly correlated with those of *MAX* (Figures S4A, S4B, and S4C). Conceivably, c-MYC is under positive selection for its oncogenic activity while MAX is not, in part due to its tumor-suppressing functions through dimerization with MXD family proteins. ^20,21^ This stoichiometric imbalance may permit, or even promote, c-MYC amyloidogenesis in malignant cancers.

### New insights into the c-MYC paradox

The “c-MYC paradox” captures the puzzling coexistence of oncogenic and tumor-suppressive activities within the same protein. Its pro-apoptotic function has traditionally been attributed solely to transcription. Our findings reveal an additional, transcription-independent mechanism: the amyloid state of c-MYC directly induces apoptosis. Thus, c-MYC can exert tumor-suppressive effects through two distinct mechanisms—transcription-dependent and transcription-independent—likely governed by its cellular abundance.

Because c-MYC expression is tightly regulated under normal conditions, disruption of this regulation is closely associated with pathogeneses, particularly malignancy. Conceptually, the amyloidogenic conversion of c-MYC may serve as a built-in failsafe mechanism: as c-MYC accumulates beyond the buffering capacity of MAX, amyloid formation promotes cytotoxicity and limits oncogenic potential.

### Implications in cancer and neurodegeneration

We previously described “tumor-suppressive amyloidogenesis” arising from widespread proteomic instability in cancer.^14^ The present findings position c-MYC as a prototypical tumor-associated amyloidogenic protein. Given the cytotoxic nature of amyloid conformers, cancer cells must neutralize them to avert tumor suppression. Our prior work showed that HSF1, the master regulator of the HSR/PSR, can physically sequester toxic AOs. ^22^ Notably, HSF1 interacts directly with c-MYC. ^15^ It is therefore plausible that HSF1 counteracts c-MYC amyloids in cancer cells, consistent with the significant positive correlation between *HSF1* and *c-MYC* expression across human cancers.^15^

Beyond cancer, our data reveal that c-MYC amyloidogenesis occurs in human AD brain tissues. In fact, elevated MYC expression has been reported in human AD brains.^23,24^ Moreover, neuronal overexpression of *c-MYC* induced neurodegeneration in mice, previously attributed to neuronal cell cycle re-entry. ^25^ Our findings, nevertheless, raise the possibility that c-MYC amyloidogenesis contributes to neurodegenerative processes—a hypothesis warranting further investigation.

## ACKNOWLEDGMENTS

This research was supported by the Intramural Research Program of the National Institutes of Health (NIH), National Cancer Institute, Center for Cancer Research (1ZIABC011767). The contributions of the NIH authors were made as part of their official duties as NIH federal employees, are in compliance with agency policy requirements, and are considered Works of the United States Government. However, the findings and conclusions presented in this paper are those of the authors and do not necessarily reflect the views of the NIH or the U.S. Department of Health and Human Services.

## Author contributions

L. L., and K-H. C. designed and conducted the experiments. C.D. conceptualized and supervised this study and analyzed the results. C.D. wrote the manuscript.

## Declaration of interests

The authors declare no competing interests.

## EXPERIMENTAL METHODS

### Cell Lines and Reagents

HeLa (female) cells and NIH3T3 (male) cells were purchased from ATCC. They were authenticated by ATCC by STR profiling. All cell cultures were maintained in DMEM supplemented with 10% HyClone bovine growth serum and 1% penicillin–streptomycin. Cells were maintained in an incubator with 5% CO2 at 37 °C. All cell lines were routinely tested for mycoplasma contamination using MycoAlert Mycoplasm Detection kits.

Recombinant proteins were all purchased commercially, including c-MYC/MAX complexes (cat#81087, Active Motif Inc.), His-HSF1 (cat#ADI-SPP-900-F, Enzo Life Sciences Inc.), GST-c-MYC (cat#M86-30G, Sino Biological US Inc.), His-c-MYC (cat#230-00580-100, RayBiotech Inc.), GST (cat#G52-30U-50, SignalChem Inc.), His-GST (cat#RPT0002, ABclonal Inc.), His-MAX (cat#81017, Active Motif Inc.), His-USF1 (cat#228-21891-2, RayBiotech Inc.), and His-OCT4 (cat#228-21139-2, RayBiotech Inc.).

All antibodies were purchased commercially, including rabbit polyclonal anti-amyloid oligomer (A11) Ab (cat#SPC-506, StressMarq Biosciences Inc.), rabbit monoclonal anti-c-MYC/N-MYC (D3N8F) Ab (cat#13987, Cell Signaling Technology), rabbit monoclonal ani-MAX (E6F6Y) Ab (cat#17471, Cell Signaling Technology), mouse monoclonal anti-HSC/HSP70 (W27) Ab (cat#sc-24, Santa Cruz Biotechnology Inc.), mouse monoclonal anti-HSP90α/β (F-8) Ab (cat#sc-13119, Santa Cruz Biotechnology Inc.), rabbit monoclonal anti-cleaved caspase 3 (Asp175) (D3E9) Ab (cat#9579, Cell Signaling Technology), goat polyclonal anti-c-MYC Ab (cat#AF3696, R&D Systems Inc.), mouse monoclonal anti-HA Ab (cat#901513, BioLegend Inc.), mouse monoclonal anti-βActin (GT5512) Ab (cat#GTX629630, GeneTex Inc.), normal rabbit IgG (cat#02-6102,ThermoFisher Scientific Inc.), Peroxidase AffiniPure Goat Anti-Rabbit IgG (H+L) (cat#111-035-144, Jackson ImmunoResearch Inc.), Peroxidase AffiniPure Goat Anti-Mouse IgG (H+L) (cat#115-035-003, Jackson ImmunoResearch Inc.), and Peroxidase IgG Fraction Monoclonal Mouse Anti-Goat IgG, light chain specific (cat#205-032-176, Jackson ImmunoResearch Inc.).

All chemicals were purchased commercially, including Thioflavin T (ThT) (cat# AC211760050, ThermoFisher Scientific Inc.), Congo red (CR) (cat#C580-25, ThermoFisher Scientific Inc.), Janus Green B (cat# AC191680050, Fisher Scientific LLC), and 10058-F4 (cat#T3048, TargetMol Chemicals Inc.).

pcDNA-GFP and pcDNA3-2xHA-MYC were gifts from Martine Roussel (Addgene plasmid#74165 and 74164, respectively). pcDNA3-2xHA-MYC^ΔC^, 2xHA-MYC^ΔC+ΔP2^, 2xHA-MYC^ΔC+ΔP12^, and 2xHA-MYC^ΔC+ΔP2+ΔP12^ were generated using the Q5 Site-Directed Mutagenesis kit (cat#E0554S, New England Biolabs Inc.). pMYC-SEAP was purchased from Clontech Laboratories (Cell signaling pathway profiling systems, cat#631910) and pCMV-Gaussia Luc was purchased from Thermo Fisher Scientific (cat#16147).

The c-MYC peptide library and alanine scan libraries were custom synthesized by GenScript USA Inc.

Paraffin sections of human liver cancer (cat#HuCAT081) and prostate cancer (cat#HuCAT371) were purchased from TissueArray.com LLC. Paraffin sections of human Alzheimer’s disease brains (cat#CSA0225P) were purchased from American MasterTech Scientific. Paraffin sections of human A2058 melanomas were prepared from xenografted tumors described previously. ^14^

### Soluble and Insoluble Fractionation

HeLa cells were treated with Congo red for 24 hours, followed by incubation at 43°C for 30 min. Cells were then trypsinized, washed, and resuspended in 1xPBS for cell counting. Equal numbers of cells were used for soluble and insoluble fractionation. The detailed procedures were described previously. ^22^

### In-Cell PLA ELISA

To avoid the interference of CR or 10058-F4 in PLA fluorescence detection, In-Cell PLA ELISA was adopted using the Duolink® In Situ Detection Reagent Brightfield (cat#DUO92012, Sigma-Aldrich). The detailed procedures were described previously.^15^ HeLa cells were seeded in 96-well culture plates (7000 cells/well) and treated with CR for 24 hr or 10058-F4 for 30 hr. The goat anti-c-MYC antibody and the rabbit anti-AO (A11) antibody were combined for the PLA, followed by incubation with Duolink® PLA anti-rabbit Plus (cat#DUO92002, Sigma-Aldrich) and anti-goat Minus probes (cat#DUO92006, Sigma-Aldrich). Of note, 1-Step Ultra TMB-ELISA Substrate Solution (cat#34029, ThermoFisher Scientific), instead of the precipitating substrate provided by the kit, was used for the substrate development.

### Immunoblotting and Immunoprecipitation

Whole-cell protein extracts were prepared in cold cell-lysis buffer (50mM Tris-Cl, pH 7.5, 120mM NaCl, 0.5% NP-40 and 1mM PMSF). Proteins were transferred to nitrocellulose membranes. Following incubation with the blocking buffer (5% non-fat milk in 1x TBS-T) for 1 hour at RT, membranes were incubated with primary antibodies (1:1,000 dilution in the blocking buffer) overnight at 4 °C. After washing with 1xTBS-T for 3 times, membranes were incubated with peroxidase-conjugated secondary antibodies (1: 5000 diluted in the blocking buffer) at RT for 1 hr. Signals were detected using SuperSignal West chemiluminescent substrates (cat#34580 or 34095, Thermo Fisher Scientific Inc.).

For the A11 IP, HeLa cells were seeded for 24 hours or transfected with plasmid DNAs for 24 hours. Following trypsinization and washing in ice-cold 1xPBS, cells were lysed in cell lysis buffer on ice for 30 min and the extracts were centrifuged at 16,813×g for 5 min at 4°C to remove insoluble fractions. Soluble cell lysates (5 mg) were incubated with either rabbit IgG or anti-amyloid oligomers (A11) antibody (5 μg) at 4°C with gentle rotation for overnight. The immune complexes were then recovered with Dynabeads™ Protein G (cat#10004D, Life Technologies Corp.) or ChIP-Grade Protein G agarose beads (cat#9007S, Cell Signaling Technology). After washing with the lysis buffer, precipitates were subjected to SDS-PAGE and immunoblotting with either goat anti-c-MYC Abs or mouse anti-HA Abs, followed by incubation with HRP-conjugated bovine anti-goat IgG (cat#805-035-180, Jackson ImmunoResearch Laboratories Inc.) or EasyBlot® anti-mouse IgG (HRP) (cat#GTX221667-01, GeneTex Inc.). Signals were detected using SuperSignal™ West Pico PLUS chemiluminescent substrates and documented with an iBright FL1000 Imaging System (ThermoFisher Scientific).

### *In Situ* Proximity Ligation Assay (PLA)

The general procedures for *in situ* PLA were described previously.^15^ For the specific MYC-A11 *in situ* PLA, the goat anti-c-MYC antibody and the rabbit anti-AO (A11) antibody were used, followed by incubation with Duolink® PLA anti-rabbit Plus and anti-goat Minus probes. Images were captured using a Zeiss LSM780 confocal microscope.

Brightfield *in situ* PLA was performed using the Duolink® In Situ Detection Reagents Brightfield. Paraffin-embedded tissue sections were deparaffinized at 60°C for 1.5 hr and rehydrated. Antigen retrieval was then performed using sodium citrate buffer (pH 6.0) in a vegetable steamer for 25 min. Following permeabilization using 0.3% Triton-X-100 in 1xPBS at RT for 10 min, endogenous peroxidase and alkaline phosphatase activities were quenched using the BLOXALL blocking solution at RT for 10 min and slides were blocked with 1x Duolink blocking solution (cat#DUO82007, Sigma-Aldrich) at 37°C for 1 hr. The tissue slides were incubated with rabbit anti-AO (A11) Abs (1:100) and goat anti-c-MYC Ab (1:100) or goat normal IgG (Jackson ImmunoResearch Labs) at RT for 3 hours, followed by incubation with anti-rabbit Plus and anti-goat Minus probes at 37°C for 1 hour, ligation at 37°C for 30 min, and amplification at 37°C for 100 min. After washing, the tissue sections were incubated with the detection reagent at RT for 1 hr, and the signals were developed with substrate at RT for 5 to 15 min and counterstained with nuclear staining solution. The slides were mounted for imaging.

### *In Vitro* Fibrillation Assay

Reactions were carried out in non-binding black 96-well microplates (cat#655900, Greiner Bio-One North America Inc.). Each well contains 10 nM recombinant proteins or 100 nM c-MYC peptides with 10 μM Thioflavin T (ThT) in 1xPBS. Plates were incubated at 37°C with gentle shaking at 220 rpm in an Eppendorf ThermoMixer C (Eppendorf North America). ThT fluorescence was measured at Ex 450 nm/Em 500 nm using a CLARIOstar microplate reader (BMG LABTECH). To correct for the differential background signals, the baseline fluorescence readings (*t_0_*) were subtracted.

### A11^+^–AO ELISA

Following *in vitro* fibrillation assays, reaction solutions in multiple wells of 96-well microplates were collected and centrifuged at 10,000 rpm for 5 min. The collected supernatants were coated on 96-well UltraCruz® ELISA microplates (cat#sc-204463, Santa Cruz Biotechnology Inc.), 100 μl per well, at 4°C overnight. Plates were washed with 1xPBS and blocked with 5% nonfat milk in 1xTBS-T at RT for 1 hour. Following blocking, diluted A11 antibody (1:1000) in the blocking buffer was added 100 μl/well and incubated at 4°C overnight with gentle shaking. After washing with 1xTBS-T, each well was incubated with 100 μl of goat anti-rabbit IgG) poly-HRP conjugates (cat#PI32260, ThermoFisher Scientific, 1:1000 dilution in the blocking buffer) at RT for 1 hour. Following washing, 100 μl of 1-Step Ultra TMB-ELISA substrates was added to each well. The plates were incubated at RT on a shaker for 2-5 min, followed by measuring the OD at 650 nm using a SpectraMax iD5 multi-mode microplate reader (Molecular Devices).

### Transmission Electron Microscopy

Recombinant proteins were first desalted and reconstituted in 1xPBS. Protein concentrations were adjusted to 0.5 μM with 1xPBS in 96-well non-binding microplates and incubated at 37°C with gentle shaking at 400 rpm in an Eppendorf ThermoMixer C (Eppendorf North America).

Following incubation for 5 days, 10 μl of each reaction solution was applied on a piece of parafilm. Then, a water-treated 400-mesh carbon-coated copper grid (cat#CF400-Cu-50, Electron Microscopy Sciences Inc.) was immersed in the droplet of sample suspensions for 1 min, and the remaining liquid was wicked away. The grids were immediately incubated with a drop of 2% uranyl acetate solution (cat#22400-2, Electron Microscopy Sciences Inc.) for 1 min. After wicking, the grids were air-dried and examined under a Hitachi HT7800 transmission electron microscope (Hitachi, Ltd. Group, Japan) operating at 120 kV.

### Transfection and c-MYC dual reporter assays

All plasmids were transfected with TurboFect™ transfection reagents (cat# R0531, Thermo Fisher Scientific). HeLa cells were co-transfected with pMYC-SEAP and pCMV-Gaussia luciferase (GLuc) reporter plasmids, along with various indicated plasmids for 24 hr. Reporter activities in culture media were measured. SEAP and GLuc activities in culture supernatants were quantitated using a NovaBright Phospha-Light EXP Assay Kit (cat# N10578, Thermo Fisher Scientific) for SEAP and a Pierce™ Gaussia Luciferase Glow Assay Kit (cat#16160, Thermo Fisher Scientific), respectively. Luminescence signals were measured by a CLARIOstar microplate reader (BMG LABTECH). SEAP activities were normalized against GLuc activities.

### Detection of Apoptosis

NIH3T3 cells were seeded in 6-well culture plates and transfected with plasmid DNAs for 24 hours. The cell culture medium was refreshed with 0.5% serum-containing DMEM for 20 hours. Cells were then collected by trypsin-EDTA digestion, washed and fixed with 4% formaldehyde at RT for 30 min. Fixed cells were subsequently permeabilized with 0.3% Triton X-100 in 1xPBS at RT for 10 min. Following wash with 1xPBS, cell pellets were resuspended and incubated with 150 µL of rabbit anti-cleaved caspase-3 Abs (cat#9661, Cell Signaling Technology, 1:200 diluted with 5% BSA in 1xPBS) at 4°C overnight. Subsequently, cells were washed and incubated with Alexa Fluor 647-conjugated goat anti-rabbit IgG (H+L) (cat# A-21245, Thermo Fisher Scientific, 1:700 diluted with 5% BSA in 1xPBS) for 1h at RT. Following washing and resuspending in 1xPBS, cells were subjected to quantification by Flow Cytometry. For detection of Alexa Fluor 647, a BD LSRFortessa™ flow cytometer (BD Biosciences, San Jose, CA) with red laser excitation at 640 nm and emission detector bandpass filter of 670/30 nm was used. Data were collected using BD FACS Diva software version 8.0.1.

## QUANTIFICATION AND STATISTICAL ANALYSIS

Statistical analyses were performed using Prism 10 (GraphPad Software). All results are expressed as mean±SD, mean±SEM, or median±IQR. The statistical significance is defined as: *p<0.05, **p<0.01; ***p<0.001; ****p<0.0001; n.s.: not significant. The *in vitro* and *ex vivo* experiments were not randomized.

## Notes

### Competing Interest Statement

The authors have declared no competing interest.

### Summary of Updates

the revised version includes updated figures and main text.

