## Supplemental Figures for "c-MYC is an aggregation-prone, amyloidogenic protein"

**LIST OF SUPPLEMENTARY MATERIALS**

The supplementary materials include 4 supplementary figures and their legends.

**Figure S1: c-MYC displays amyloid-like properties under pathological conditions, related to Figure 1.**

**Figure S2: c-MYC is intrinsically amyloidogenic, related to Figure 2.**

**Figure S3: The amyloidogenic P2 and P12 regions are disordered, related to Figure 3.**

**Figure S4: The amyloidogenesis of c-MYC contributes to its intrinsic tumor suppressor activity, related to Figure 4 and Discussion.**

### Figure S1

**A**

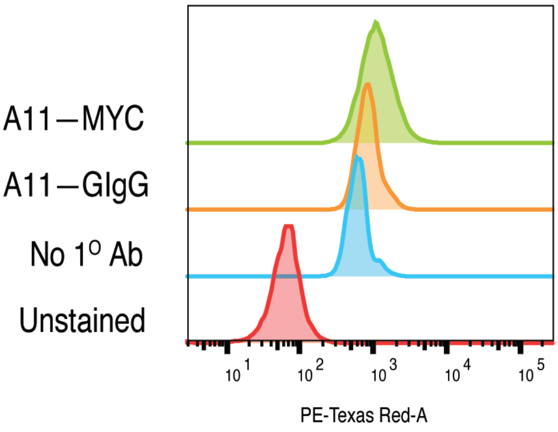

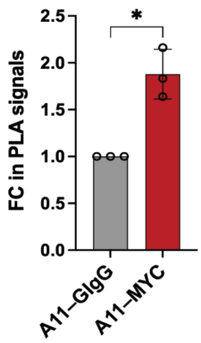

**B**

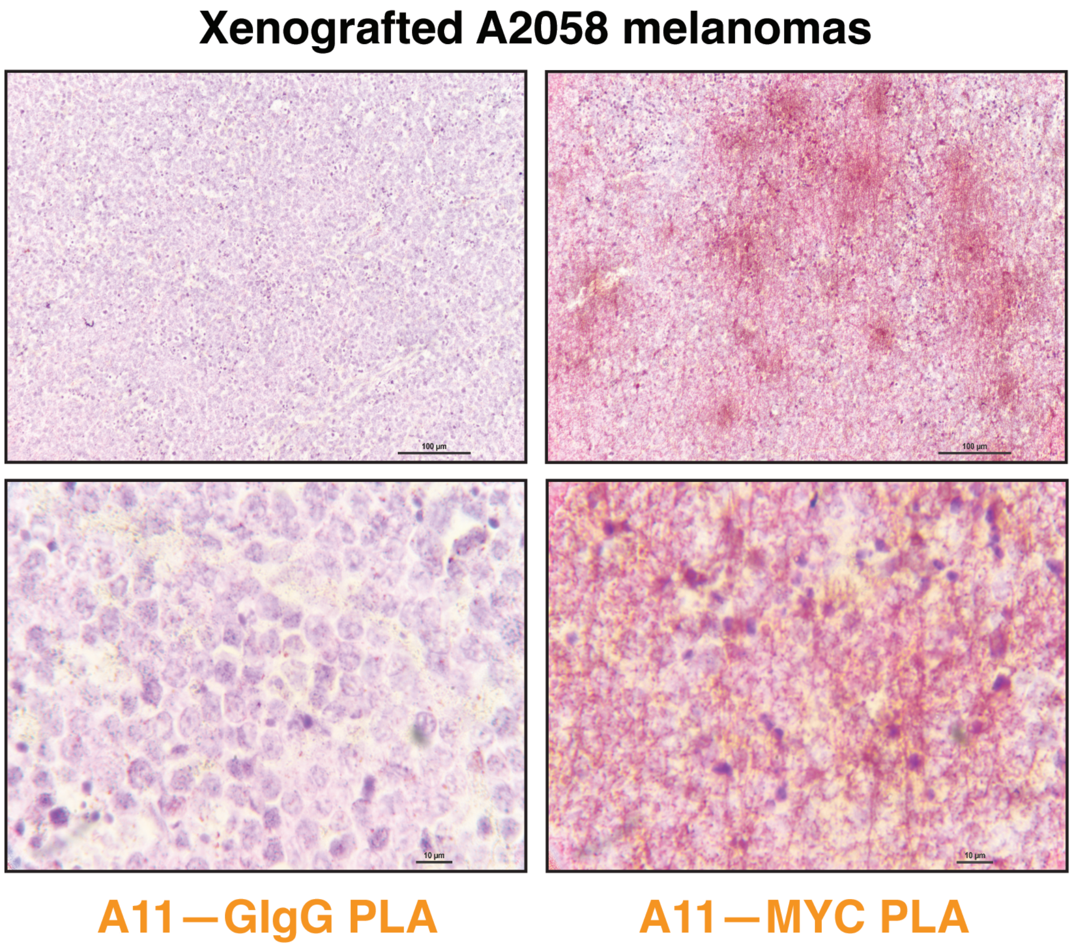

**C**

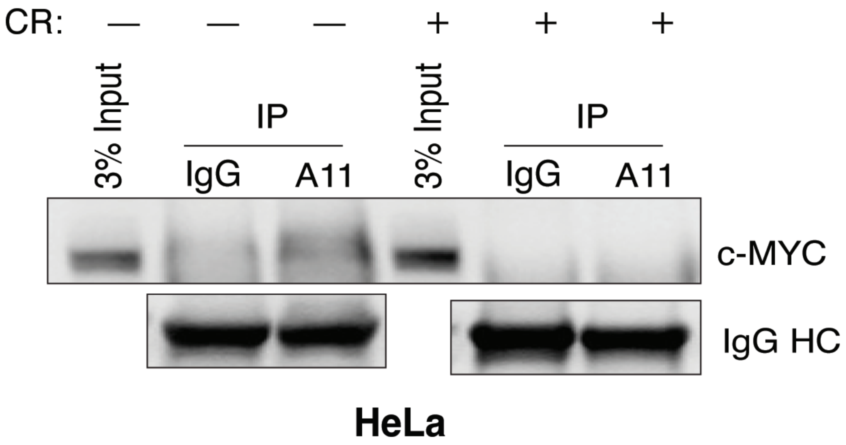

#### SUPPLEMENTAL FIGURE LEGENDS

**Figure S1: c-MYC displays amyloid-like properties under pathological conditions.** (A) Quantitation of endogenous c-MYC recognized by both the goat anti-c-MYC Ab and the rabbit anti-AO (A11) Ab through flow PLA (mean ± SD, n=3 independent experiments, Two-tailed Student's *t* test). (B) Detection of endogenous c-MYC proteins in xenografted human A2058 melanomas by brightfield PLA using both the goat anti-c-MYC Ab and the rabbit anti-AO (A11) Ab. Scale bars: 100 μm for low magnification and 10 μm for high magnification. (C) Immunoprecipitation of endogenous c-MYC proteins by A11 antibodies in HeLa cells treated with 30 μM CR.

### Figure S2

A

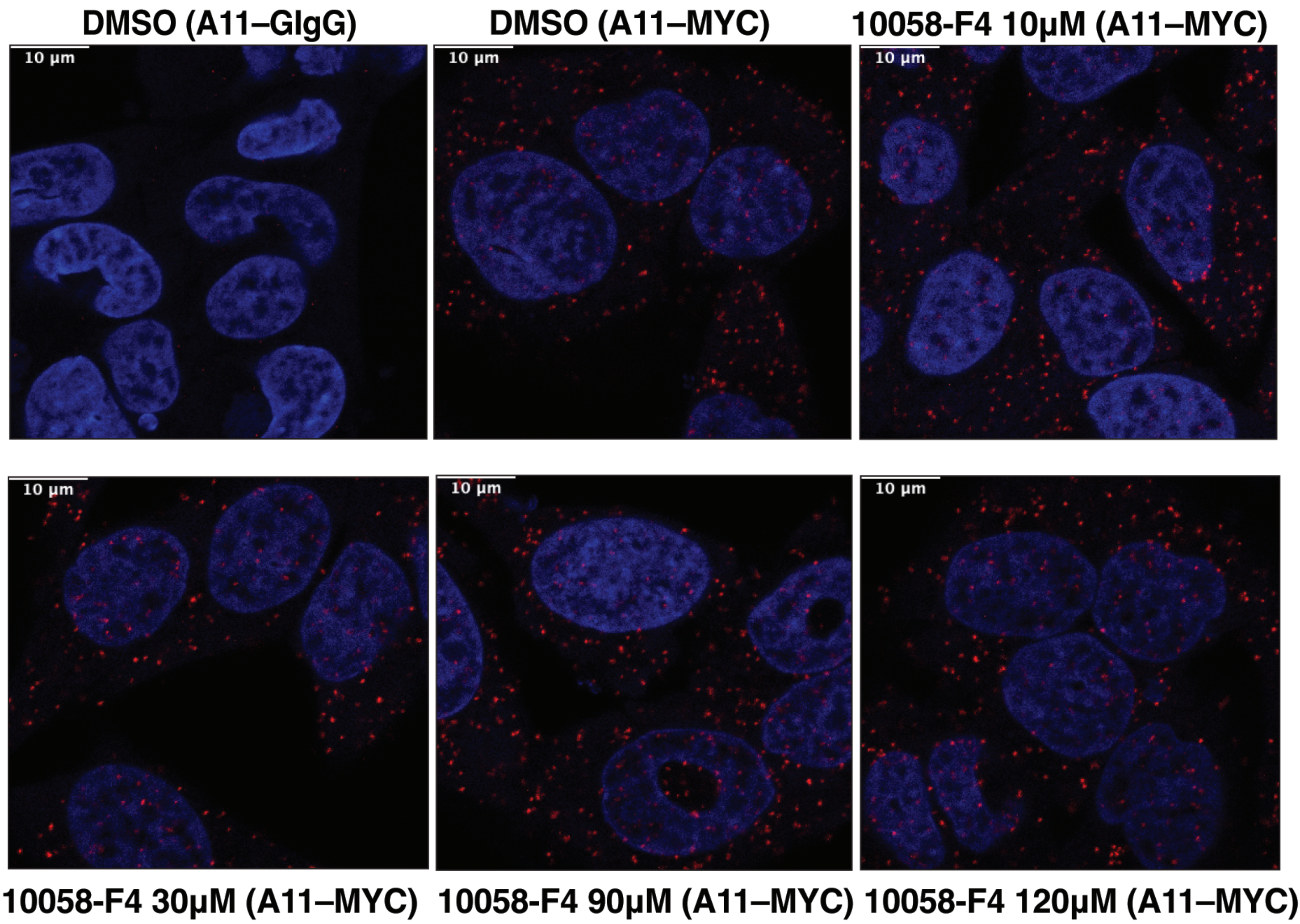
DMSO (A11–GIgG)
DMSO (A11–MYC)
10058-F4 10μM (A11–MYC)
10 μm
10058-F4 30μM (A11–MYC)
10058-F4 90μM (A11–MYC)
10058-F4 120μM (A11–MYC)

B

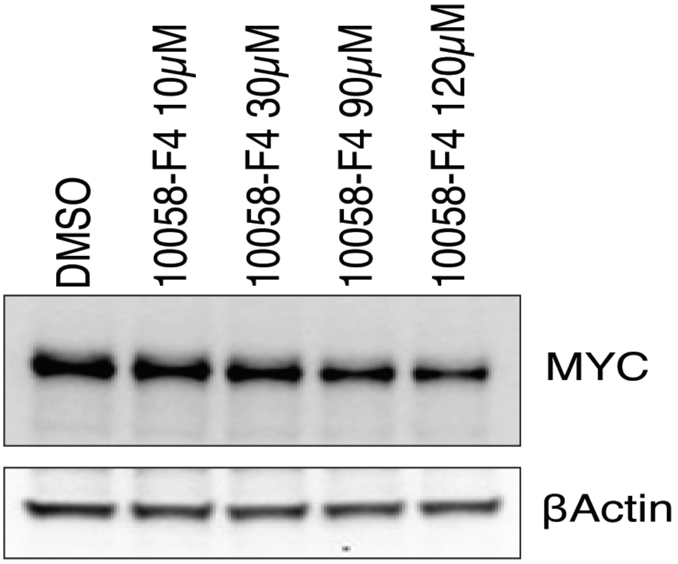
DMSO
10058-F4 10μM
10058-F4 30μM
10058-F4 90μM
10058-F4 120μM
MYC
βActin

**Figure S2: c-MYC is intrinsically amyloidogenic.** (A) Visualization of endogenous c-MYC AOs in HeLa cells treated with 10058-F4 by fluorescent PLA (representative images of three independent experiments). Scale bars: 10 μm. (B) Detection of c-MYC proteins by immunoblotting in HeLa cells treated with 10058-F4 (representative images of three independent experiments).

### Figure S3

**A**

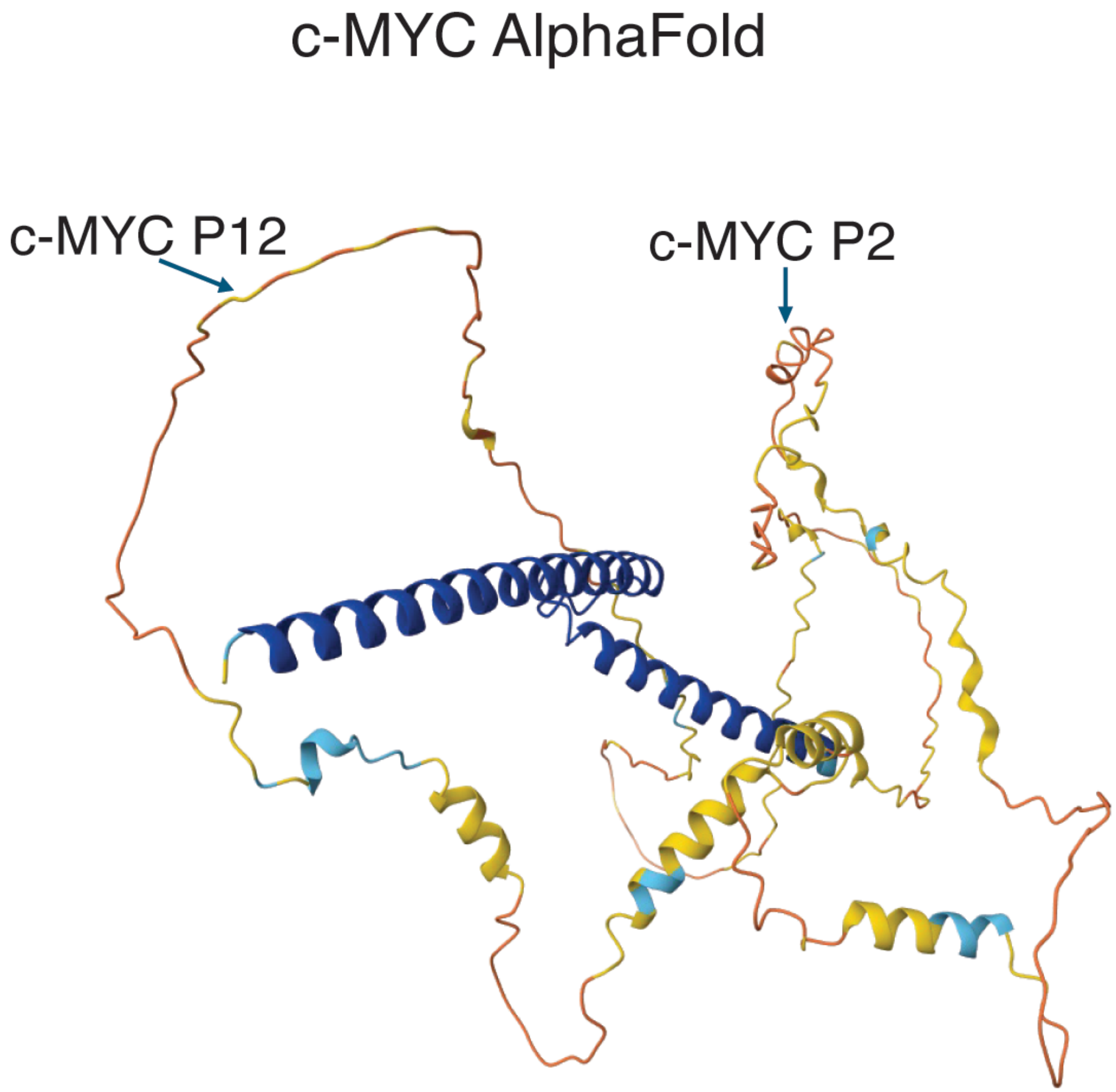

**Figure S3: The amyloidogenic P2 and P12 regions are disordered.** (A) Predicted 3D structures of human c-MYC proteins by AlphaFold, with the P2 and P12 regions highlighted.

### Figure S4

A

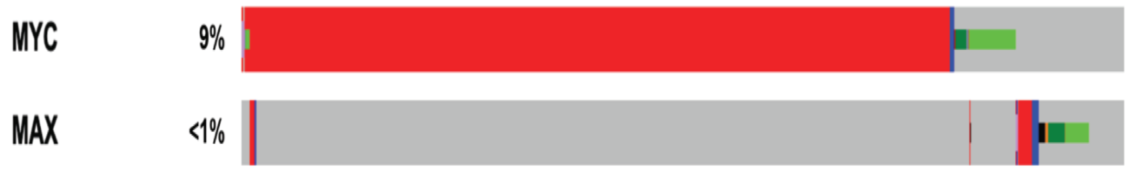

B

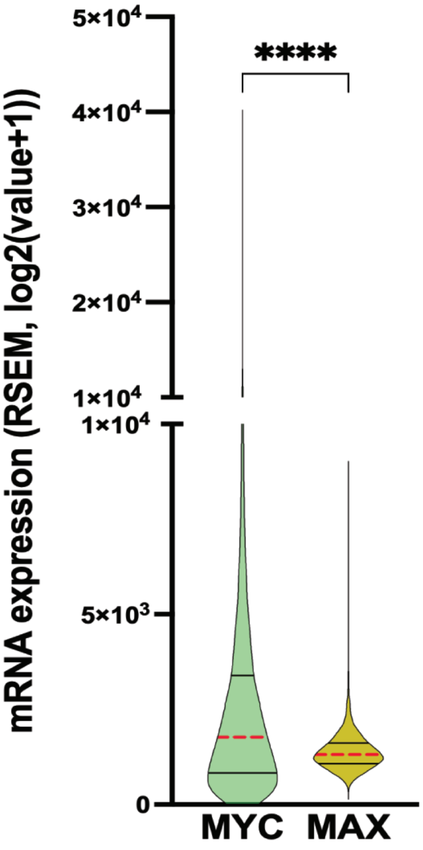

C

| Gene | Spearman correlation | *P* value | Pearson correlation | *P* value |
|---|---|---|---|---|
| *MAX* | 0.06 | 1.59e-09 | 0.06 | 6.17e-10 |
| *HSF1* | 0.22 | 1.52e-108 | 0.23 | 9.09e-125 |
| *MXD1* | 0.16 | 3.94e-59 | 0.13 | 2.33e-36 |
| *MXD3* | -0.07 | 3.70e-13 | -0.07 | 2.72e-11 |
| *MXI1* | 0.17 | 7.70e-69 | 0.17 | 2.32e-63 |
| *MNT* | 0.13 | 1.29e-37 | 0.11 | 9.82e-29 |
| *MIZ1* | -0.09 | 8.75e-20 | -0.07 | 1.47e-11 |

**Figure S4: The amyloidogenesis of c-MYC contributes to its intrinsic tumor suppressor activity. (**A) OncoPrint graph derived from TCGA showing the genomic alterations of *c-MYC* and *MAX* genes in human pan-cancer tissues. (B) Violin plots showing the mRNA expression of *c-MYC* and *MAX* in human tumor samples (median ± IQR, n=10071, Wilcoxon matched-pairs signed rank test). Data are derived from TCGA pan-cancer atlas. RSEM is batched normalized from Illumina HiSeq_RNASeqV2. (C) Gene expression correlation between *c-MYC* and key components of its network in human pan-cancer tissues. Data are derived from TCGA. Genes in red are activators of c-MYC-mediated transcription; genes in black are repressors of c-MYC-mediated transcription.
